# Only *SF3B1* Mutation involving K700E (And Not Other Codons), Independently Predicts Overall Survival in Myelodysplastic Syndromes

**DOI:** 10.1101/2020.09.04.283598

**Authors:** Rashmi Kanagal-Shamanna, Guillermo Montalban-Bravo, Koji Sasaki, Elias Jabbour, Carlos Bueso-Ramos, Yue Wei, Kelly Chien, Tapan Kadia, Farhad Ravandi, Gautam Borthakur, Kelly A. Soltysiak, Naval Daver, Faezeh Darbaniyan, Mark Routbort, Keyur Patel, L. Jeffrey Medeiros, Sherry Pierce, Hagop Kantarjian, Guillermo Garcia-Manero

## Abstract

**Background:** *SF3B1* mutations (*SF3B1*^*mut*^) in myelodysplastic syndromes (MDS) frequently involve codon K700E and have a favorable prognosis. The prognostic effect of non-K700E *SF3B1*^*mut*^ is uncertain.

**Methods:** We analyzed the clinical-pathologic features and outcomes of a single-institutional series of 94 *SF3B1*^*mut*^ and 415 *SF3B1*^*wt*^ newly diagnosed untreated MDS patients and explored the differences between K700E and non-K700E subgroups.

**Findings:** Ninety-four (19%) patients had *SF3B1*^*mut*^: median age, 74 years. Fifty-five (60%) patients carried K700E. Recurrent non-K700E mutations (39, 40%) included R625, H662 and K666. Compared to *SF3B1*^*mut*^ K700E, non-K700E patients had a lower median ANC (1·8 vs. 2·4, p=0·005) and were frequently “high” R-IPSS (revised International Prognostic Scoring System) [7(19%) vs. 2(4%), p=0·031]. Non-K700E MDS frequently associated with *RUNX1* (26% vs. 7%, p=0·012) and exclusively with *BCOR, IDH2*, and *SRSF2* mutations. There was no significant difference in karyotype or *SF3B1* variant allele frequency. Most (∼80%) were treated with hypomethylating agents. *SF3B1*^*mut*^ had superior overall survival (OS) than *SF3B1*^*wt*^ in all MDS categories [not-reached vs. 25·2 months, p=0·0003], low-grade MDS, and MDS with ring sideroblasts (MDS-RS). Compared to *SF3B1*^*wt*^, *SF3B1*^*mut*^ K700E had superior outcomes in all MDS categories (25 months vs. not-reached, p=0·0001), low-grade MDS, and MDS-RS, but no significant difference was seen with non-K700E. By multivariate analysis, absence of *SF3B1*^*mut*^ K700E (not non-K700E) independently associated with prognosis.

**Interpretation:** *SF3B1*^*mut*^ MDS show distinct clinical and mutational profiles, with K700E showing a significantly better OS compared to non-K700E mutations and *SF3B1*^*wt*^. Our study highlights the importance of *SF3B1* mutation type in MDS risk assessment.

**Data Sharing Statement:** The datasets generated during and/or analyzed during the current study are not publicly available due to patient privacy concerns but are available from the corresponding author on reasonable request.

**Research in Context:** *Evidence before this study:* We designed this study based on the collective evidence from a systematic search of the literature for outcomes of patients MDS with *SF3B1* mutations (*SF3B1*^*mut*^) from January 2013 to June 2020. Both the International Working Group for the Prognosis of MDS (IWG-PM) proposal and 2016 revisions to the World Health Organization (WHO) Classification of Myelodysplastic Syndromes recognize *SF3B1*^*mut*^ MDS with <5% blasts (or ring sideroblasts >5% for WHO) as a distinct sub-category, in the absence of other unfavorable features. This was largely based on favorable prognostic outcomes, a distinct gene expression profile, and association with ring sideroblasts. However, the natural history of *SF3B1*^*mut*^ MDS is heterogeneous. A high proportion of *SF3B1* mutations occur within codon K700, leading to large-scale mRNA downregulation due to branch point recognition error, while the rest occur outside of this codon. The downstream functional effects of *SF3B1* mutations outside of the K700 codon are unclear. The clinical course of *SF3B1*^*mut*^ MDS patients likely depends on the type of *SF3B1* mutation and other features such as variant allele frequency, concomitant gene mutations, and karyotype. Until now, the effects of the different types of *SF3B1* mutations were largely unknown.

*Added value of this study:* In this study, we report distinctive clinicopathologic characteristics and outcomes of MDS patients with *SF3B1* mutations segregated based on mutation type: K700E vs. non-K700E. We show that ∼40% of *SF3B1* mutated MDS patients have non-K700E mutations. Non-K700E *SF3B1*^*mut*^ MDS have distinct clinico-pathologic features, such as lower ANC and frequent association with mutations in *RUNX1, BCOR, IDH2*, and *SRSF2*. There was no significant difference in karyotype or *SF3B1* variant allele frequency. Importantly, K700E *SF3B1*^*mut*^ MDS had superior overall survival compared to *SF3B1*^*wt*^, in all MDS, low-grade MDS, and MDS with ring sideroblasts, but no significant difference was seen with non-K700E. By multivariate analysis, absence of *SF3B1*^*mut*^ K700E, but not non-K700E, independently associated with prognosis.

*Implications of all the available evidence:* To the best of our knowledge, this is the first study to report these findings from a single-institutional series of MDS primarily treated with hypomethylating agents. Our study highlights the importance of determining the *SF3B1* mutation type in MDS risk assessment. These findings are important in light of the recent FDA approval of luspatercept based on the results of the MEDALIST trial that suggested sustained hematological responses in *SF3B1*^*mut*^ MDS patients.

## INTRODUCTION

*SF3B1* is one of the most frequently mutated genes in MDS.^1^ *SF3B1* mutations are detected in a third of MDS patients and are present in up to 66% of MDS sub-categories with increased ring sideroblasts (RS).^2-4^ Prior studies have indicated that the presence of *SF3B1* mutation (*SF3B1*^*mut*^) was an independently favorable prognostic factor for survival in MDS.^2,5-8^ Based on the favorable outcome, a distinct gene expression profile, and association with ring sideroblasts (RS), the 2016 revisions to the World Health Organization (WHO) Classification of Myelodysplastic Syndromes recognized MDS-RS with single lineage (MDS-RS-SLD) and multilineage dysplasia (MDS-RS-MLD) as distinct sub-categories, and recommended testing for *SF3B1* mutation in MDS patients with 5 and 15% ring sideroblasts in bone marrow (BM) aspirates.^9^ We have previously shown that the survival of MDS-RS-MLD patients (a majority of which had *SF3B1* mutations) was significantly better compared to those with MDS-MLD without RS.^10^ Based on recent data suggesting sustained hematological responses in *SF3B1*^*mut*^ MDS patients treated with luspatercept,^11^ the International Working Group for the Prognosis of MDS (IWG-PM) has proposed that *SF3B1*^*mut*^ MDS be considered a distinct entity with a favorable prognosis in the absence of >5% BM blasts or ≥1% peripheral blood (PB) blasts, absence of del(5q), monosomy 7, inv(3), abnormal 3q26 or complex karyotype (CK), mutations in *RUNX1* and *EZH2*, and findings suggestive of other WHO-defined entities.^6^

Despite this body of information, the natural history of *SF3B1* mutated MDS is heterogeneous. A high proportion of *SF3B1* mutations occur within codon 700, causing a branch point recognition error of a cryptic splice site followed by nonsense decay due to aberrant splice junctions, ultimately resulting in large-scale downregulation of mRNA.^12-14^ The remainder of *SF3B1* mutations occur outside of this hotspot and the downstream functional effects are not clear. Dalton, *et al* has recently suggested that *SF3B1* K666N mutations had distinctive RNA splicing profiles and were associated with distinct clinico-pathologic features and worse outcomes. The authors suggested that these mutations may need aggressive management, even in the setting of a lower IPSS-R category.^15^ Beyond the *SF3B1* mutation type, the clinical course can be potentially altered by other parameters, such as the variant allele frequency (VAF), presence of concomitant gene mutations and karyotype.

In this study, we investigated the clinico-pathologic and genetic features and outcomes in a single-institutional series of 94 *SF3B1*^*mut*^ MDS patients and compared them to 415 with wild-type MDS (*SF3B1*^wt^). We also explored the differences between the *SF3B1* K700E and non-K700E mutated MDS subgroups, and demonstrated distinct clinical and mutational profiles, with the K700E mutated subgroup showing a significantly better overall survival (OS) compared to the non-K700E subgroup. Further, only the *SF3B1* K700E mutation subtype independently predicted for better OS in MDS.

## MATERIALS AND METHODS

### Patients and samples

We selected BM aspirates from all newly diagnosed previously untreated MDS patients who underwent 81-gene panel next-generation sequencing (NGS) at The University of Texas MD Anderson Cancer Center (MDACC) from 2017-2019. Diagnoses were confirmed by BM examination and sub-classified using the 2016 WHO criteria.^16^ At least 20 metaphases were evaluated, and interpreted according to the 2016 International System for Human Cytogenetic Nomenclature. Risk stratification was done using the IPSS-R^17^ for patients with MDS. The study was approved by the MDACC Institutional Review Board and all samples were collected following institutional guidelines with informed consent in accord with the Declaration of Helsinki.

### Targeted next-generation sequencing

All patient samples underwent comprehensive NGS-based mutation analysis using our 81-gene panel comprising the hotspots and whole coding regions of myeloid leukemia-related genes in a CLIA-certified Molecular Diagnostics Laboratory, as previously described. ^18^ Briefly, genomic DNA was extracted from fresh BM aspirates using standard techniques, followed by library preparation and amplicon-based targeted NGS on a MiSeq sequencer. Sequences were aligned to the GRCh37/hg19 reference genome. With a minimum of 250X bidirectional coverage, a minimum variant call of 2% was considered as the limit of detection. The somatic nature of the variants was inferred based on the information in online SNP databases (*i*.*e*., Exome Aggregation Consortium [ExAC], dbSNP 137/138, 1000 Genomes) and the literature. *FLT3*-ITD mutations were assessed by PCR-based capillary electrophoresis.

### Statistical analysis

OS was calculated as the time from diagnosis to death or last follow-up date. Patients alive at their last follow-up were censored on that date. The Kaplan-Meier product limit method was used to estimate the median OS for each clinical/demographic factor. Univariate Cox proportional hazards regression analysis was used to identify any association with each of the variables and survival outcomes followed by multivariate analysis. Response assessment was performed following 2006 IWG criteria.^19^

## RESULTS

### Patient characteristics

A total of 509 treatment naïve MDS patients presented to our institution between 2017 and 2019, of which 94 (18·5%) had *SF3B1* mutations. None of the patients had received prior therapy, including erythroid stimulating agents. The clinical characteristics of patients analyzed are shown in **Table 1**. The *SF3B1*^*mut*^ MDS patients included 59 men and 35 women, with a median age of 74 (range, 39-92) years. The distribution of IPSS-R scores for this cohort were as follows: 14 (14·9%) very low; 37 (39·4%) low; 13 (13·8%) intermediate; 9 (9·6%) high; and 12 (12·8%) very high. Seventeen (18%) patients had received prior chemotherapy and/or radiation for an unrelated malignancy (therapy-related MDS, t-MDS). Among the remaining patients, 2016 WHO sub-classifications were as follows: 22 (23%) with MDS with single lineage dysplasia and ring sideroblasts (MDS-RS-SLD); 30 (32%) with MDS with ring sideroblasts and multilineage dysplasia (MDS-MLD-RS); 5 (5%) with MDS-MLD; 31 (33%) with MDS with excess blasts; 4 (4%) with MDS and isolated del(5q); and 2 (2%) with MDS-unclassifiable (MDS-U). Treatment data was available for 313 patients: 248 (79%) patients received hypomethylating agents (HMAs) alone or in combination with other agents and 35 (11·2%) received chemotherapy-based regimens.

**Table 1.**
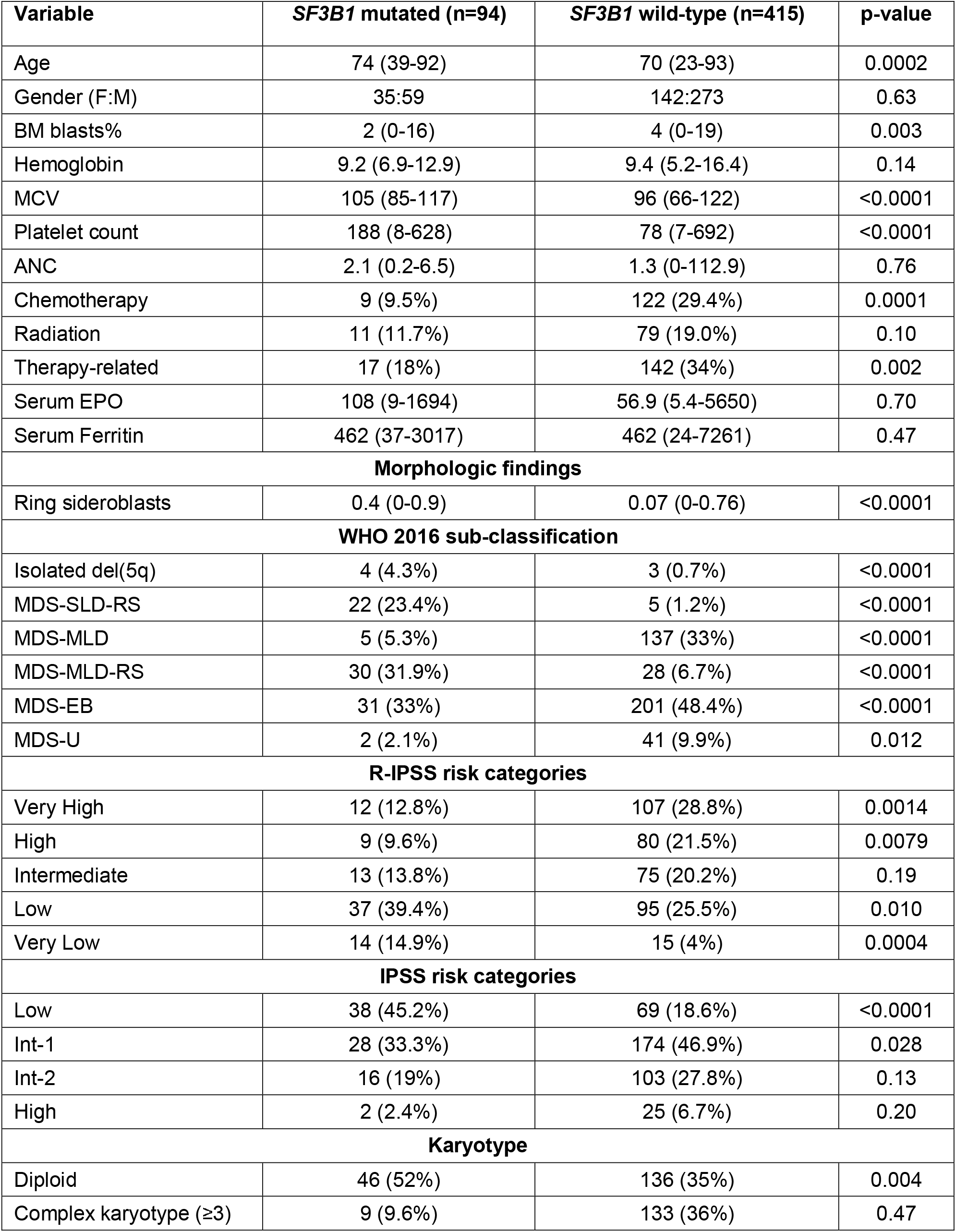
Baseline characteristics of the study group: all newly diagnosed previously untreated MDS samples that underwent mutation analysis using 81-gene next-generation sequencing panel.

| Variable | *SF3B1* mutated (n=94) | *SF3B1* wild-type (n=415) | p-value |
| --- | --- | --- | --- |
| Age | 74 (39-92) | 70 (23-93) | 0.0002 |
| Gender (F:M) | 35:59 | 142:273 | 0.63 |
| BM blasts% | 2 (0-16) | 4 (0-19) | 0.003 |
| Hemoglobin | 9.2 (6.9-12.9) | 9.4 (5.2-16.4) | 0.14 |
| MCV | 105 (85-117) | 96 (66-122) | <0.0001 |
| Platelet count | 188 (8-628) | 78 (7-692) | <0.0001 |
| ANC | 2.1 (0.2-6.5) | 1.3 (0-112.9) | 0.76 |
| Chemotherapy | 9 (9.5%) | 122 (29.4%) | 0.0001 |
| Radiation | 11 (11.7%) | 79 (19.0%) | 0.10 |
| Therapy-related | 17 (18%) | 142 (34%) | 0.002 |
| Serum EPO | 108 (9-1694) | 56.9 (5.4-5650) | 0.70 |
| Serum Ferritin | 462 (37-3017) | 462 (24-7261) | 0.47 |
| **Morphologic findings** |  |  |  |
| Ring sideroblasts | 0.4 (0-0.9) | 0.07 (0-0.76) | <0.0001 |
| **WHO 2016 sub-classification** |  |  |  |
| Isolated del(5q) | 4 (4.3%) | 3 (0.7%) | <0.0001 |
| MDS-SLD-RS | 22 (23.4%) | 5 (1.2%) | <0.0001 |
| MDS-MLD | 5 (5.3%) | 137 (33%) | <0.0001 |
| MDS-MLD-RS | 30 (31.9%) | 28 (6.7%) | <0.0001 |
| MDS-EB | 31 (33%) | 201 (48.4%) | <0.0001 |
| MDS-U | 2 (2.1%) | 41 (9.9%) | 0.012 |
| **R-IPSS risk categories** |  |  |  |
| Very High | 12 (12.8%) | 107 (28.8%) | 0.0014 |
| High | 9 (9.6%) | 80 (21.5%) | 0.0079 |
| Intermediate | 13 (13.8%) | 75 (20.2%) | 0.19 |
| Low | 37 (39.4%) | 95 (25.5%) | 0.010 |
| Very Low | 14 (14.9%) | 15 (4%) | 0.0004 |
| **IPSS risk categories** |  |  |  |
| Low | 38 (45.2%) | 69 (18.6%) | <0.0001 |
| Int-1 | 28 (33.3%) | 174 (46.9%) | 0.028 |
| Int-2 | 16 (19%) | 103 (27.8%) | 0.13 |
| High | 2 (2.4%) | 25 (6.7%) | 0.20 |
| **Karyotype** |  |  |  |
| Diploid | 46 (52%) | 136 (35%) | 0.004 |
| Complex karyotype ($\geq 3$) | 9 (9.6%) | 133 (36%) | 0.47 |

### Clinical and morphologic findings of *SF3B1*^*mut*^ MDS

Compared to *SF3B1*^*w*t^, *SF3B1*^*mut*^ MDS patients had a significantly higher median age (74 vs. 70, p=0·0008), higher median mean corpuscular volume (MCV) (105 vs. 96, p<0·0001), higher median platelet count (188 vs. 78, p<0·0001), and lower BM blast percentage (median, 2 vs. 4, p=0·003). There was no significant difference in median serum erythropoietin levels (108 vs. 57). Compared to *SF3B1*^*wt*^, *SF3B1*^*mut*^ MDS patients were more likely to be in the IPSS-R very-low (4% vs. 14·9%, p=0·0004) and low (25·5% vs. 39·4%, p=0·010) categories and less likely to be in the IPSS-R-high (21.5% vs. 9·6%, p=0·008) and very-high (28·8% vs. 12·8%, p=0·001) categories. The median percentage of ring sideroblasts was higher in patients with *SF3B1*^*mut*^ MDS (40% vs. 7%, p<0·0001) compared to the those with *SF3B1*^*wt*^; accordingly, the *SF3B1*^*mut*^ MDS group was significantly enriched in WHO categories with ring sideroblasts [MDS-SLD-RS (23·4% vs. 1·2%, p<0·0001) and MDS-RS-MLD (31·.9% vs. 6·7%, p<0·0001)] and MDS with isolated del(5q) (4·3% vs. 0·7%, p<0·0001), while they were underrepresented in MDS-MLD (5·3% vs. 33%, p<0·0001), MDS-EB (33% vs. 48·4%, p<0·0001), and MDS-U (2·1% vs. 9·9%, p=0·012).

*SF3B1*^*mut*^ MDS patients were less likely to be therapy-related [17 (18%) vs. 142 (34%); p=0·002]. However, within 24 t-MDS with >15% RS and <5% blasts, the most frequent mutation observed was in *SF3B1* (n=13, 54%), 4 of which had concurrent *TP53* mutations/CK. Six (25%) patients had *TP53* mutations/CK without *SF3B1* mutations, and 5 (20.8%) had neither *SF3B1* nor *TP53* mutations/CK.

### Mutational landscape of *SF3B1*^*mut*^ MDS

BM aspirates from all patients underwent NGS analysis with an 81-gene panel at the time of diagnosis. The most frequent *SF3B1* mutation, noted in ∼60% of all patients, was the hotspot K700E. Among the remaining mutations observed, the most frequent involved the codons H662, K666, and R625, seen in 8 patients each (**Figure 1 A**). The mutational landscape is depicted in **Figure 1 B**. Only a third of the patients had an *SF3B1* mutation as the sole driver of MDS, while the majority had concomitant mutations. The genes mutated in >10% of patients in decreasing order of frequency included *TET2* (25%), *DNMT3A* (21%), *RUNX1* (15%), *TP53* (10%), *ASXL1* (7%), *BCOR* (4%), *IDH1/2* (4%), *SRSF2* (3%), *NRAS* (3%), and *EZH2* (3%). Conventional cytogenetic analysis showed a higher frequency of normal karyotype and lower frequency of CK in *SF3B1*^*mut*^ compared to *SF3B1*^*wt*^ cases. Three *SF3B1*^*mut*^ patients had *EVI1* gene rearrangements, but the frequency was not significantly different compared to the *SF3B1*^*wt*^ group. Three *SF3B1*^*mut*^ patients had t(1;3)(p36;q21) RPN1/PRDM16, which was absent in the *SF3B1*^*wt*^ cohort.

**Figure 1.**
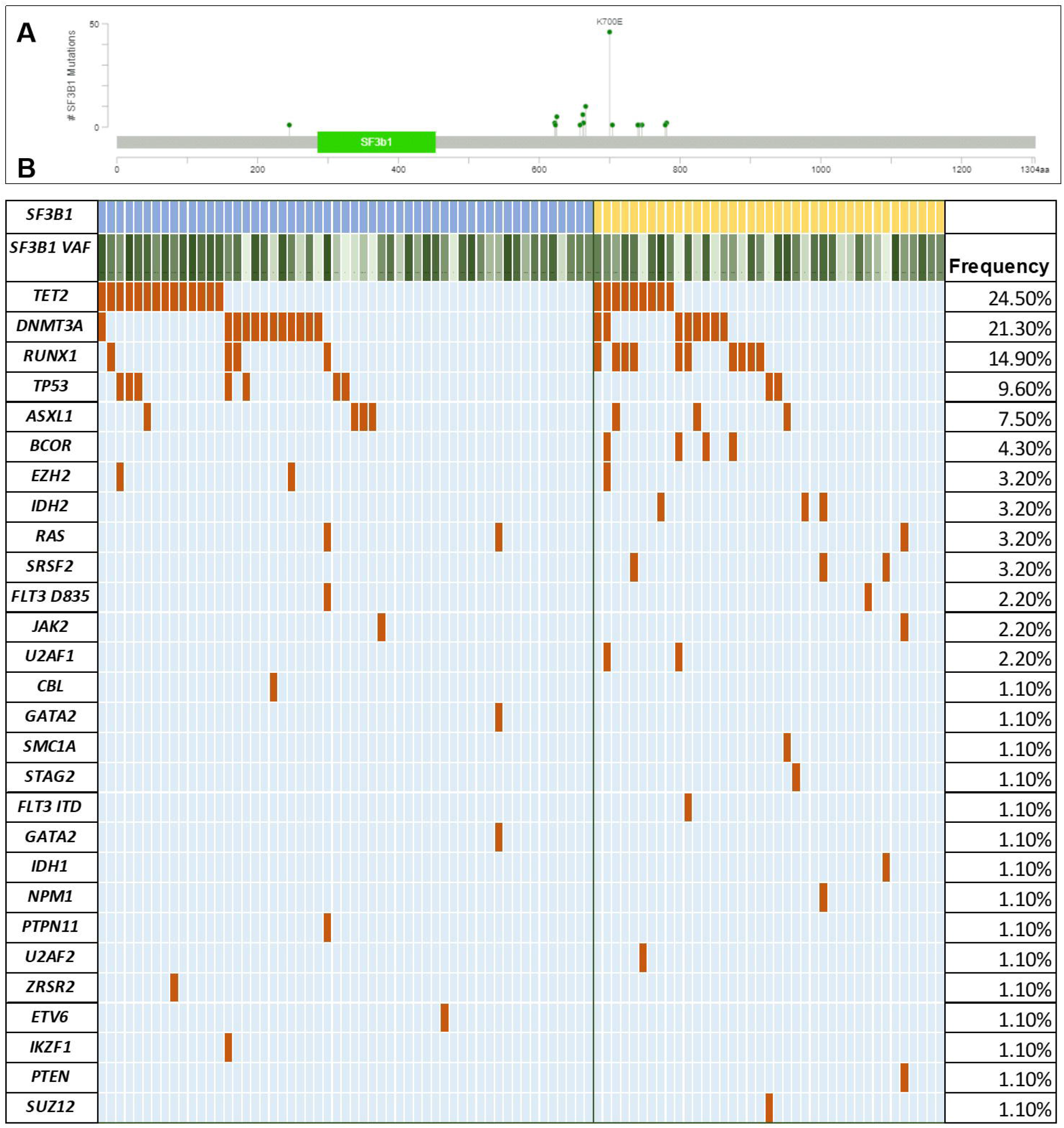
Mutational characteristics. (A) Spectrum of *SF3B1* mutations within the coding region of the gene seen in the study group generated using MutationMapper from cbioportal. About ∼60% of mutations involved the hotspot K700E, while ∼40% involved the non-K700E regions. (B) Mutational landscape of *SF3B1* K700E mutated MDS. Comutation plot of 94 *SF3B1* mutated MDS patients showing nonsynonymous gene mutations (left), separated by the mutation subtype: K700E (blue) and non-K700E (yellow). Each column represents a patient, and mutations are represented by color. The frequency of gene mutations is shown on the right. Color code: Maroon indicates mutation while blue indicates wild-type. Variant allele frequency of *SF3B1* is represented as a color gradient.

### Comparison between *SF3B1* K700E and non-K700E mutated MDS

We compared the clinico-pathologic features of 55 K700E vs. 39 non-K700E treatment naïve *SF3B1*^*mut*^ MDS patients (**Table 2**). MDS with *SF3B1* K700E mutations had a higher percentage of ring sideroblasts (median 50% vs. 34%; p=0·038), higher ANC (2·4 vs. 1·8, p=0·005), and a trend of higher platelet count (196 vs. 124, p=0·05). Among IPSS-R categories, *SF3B1*^*mut*^ MDS patients with non-K700E mutations had a significantly higher representation in IPSS-R high category [7(19%) vs. 2(4%), p=0·031], but the distribution was not significantly different among other categories. Likewise, within 2016 WHO categories, *SF3B1*^*mut*^ MDS patients with K700E mutations were less likely to be classified as MDS-EB than non-K700E *SF3B1* mutated patients (22% vs. 49%, p=0·008). All 4 patients that fit the criteria for MDS with isolated del(5q) had K700E mutations.

**Table 2.**
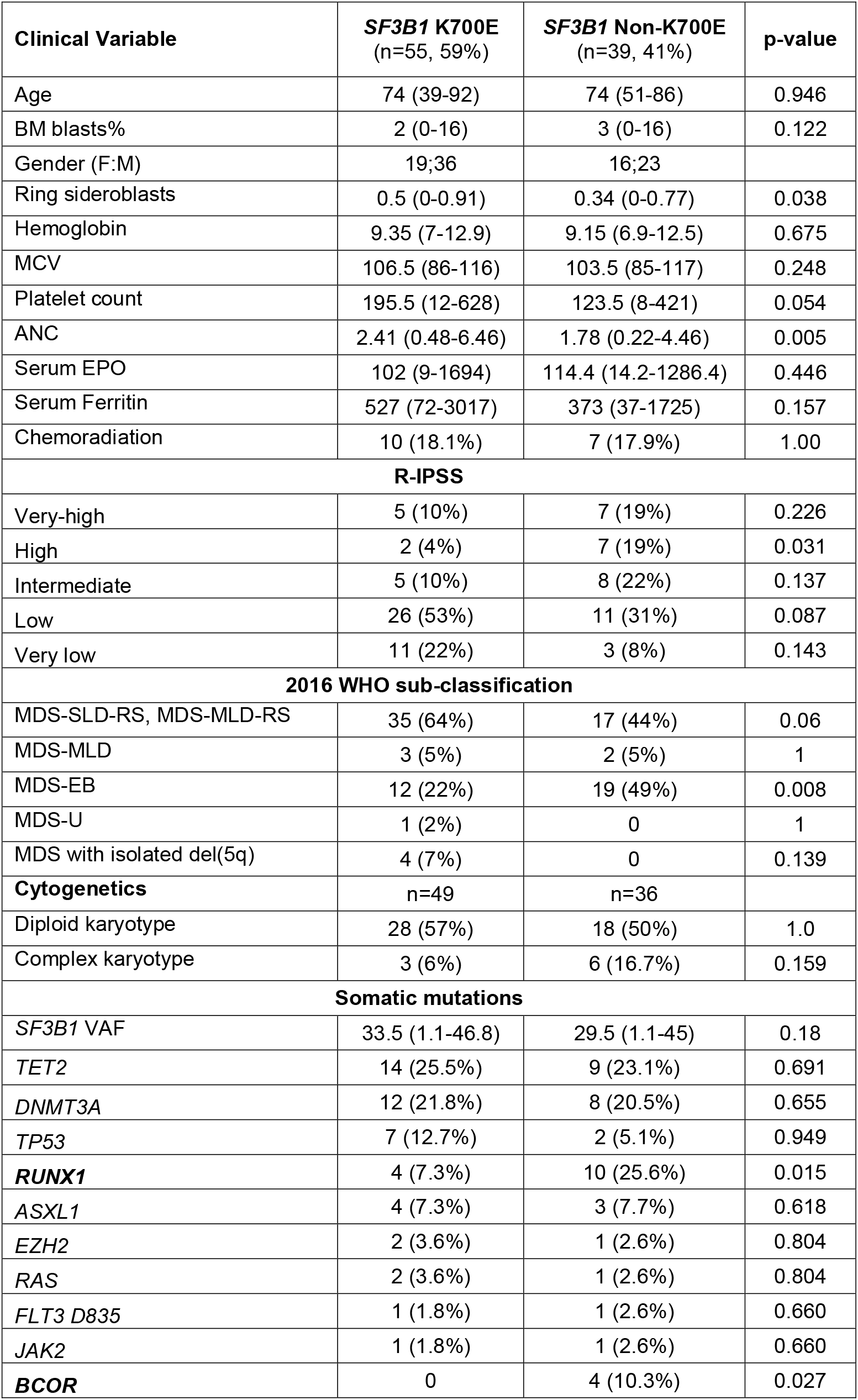
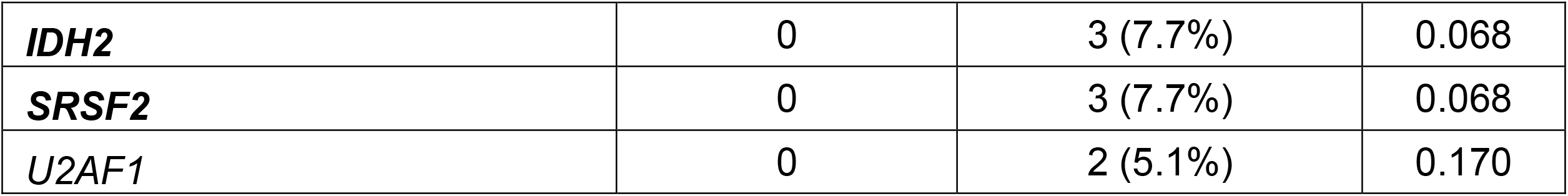
Comparison of baseline characteristics of all newly diagnosed untreated *SF3B1* mutated MDS segregated by the mutation type, K700E vs. non-K700E.

There was no difference in the median *SF3B1* variant allele frequency (VAF) between the 2 groups. The frequency of *RUNX1* mutation was significantly higher in non-K700E cases (26% vs. 7·3%, p=0·012), and mutations in *BCOR* (p=0·02*), IDH2* (p=0·07), and *SRSF2* (p=0·07) were exclusively observed in non-K700E cases (**Figure 2**). No significant differences were noted in the frequencies of *TP53* or clonal hematopoiesis-associated mutations, including *DNMT3A, ASXL1*, and *TET2*.

**Figure 2.**
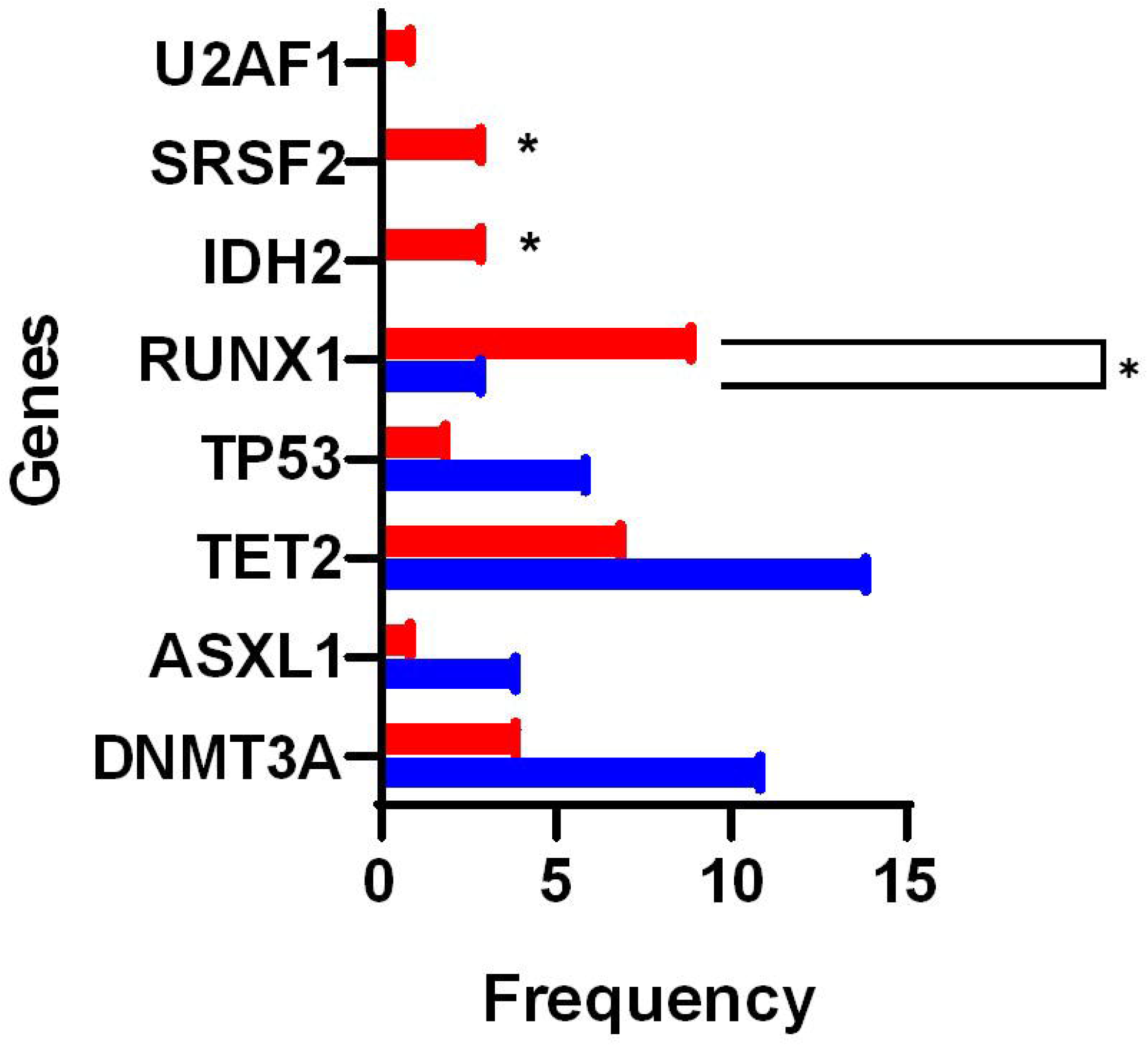
Within *SF3B1* mutated MDS, the K700E mutant group has distinct clinico-pathologic and genetic characteristics compared to non-K700E mutant MDS. *SF3B1* non-K700E mutated patients (red) had a significantly higher frequency of mutations in *RUNX1*, and a trend to higher frequencies of mutations in *SRSF2* and *IDH2* genes compared to *SF3B1* K700E mutated MDS patients.

There were no significant differences in normal vs. complex karyotype. When cytogenetic aberrations were classified using the comprehensive cytogenetic scoring system (CCSS; scores from 0-5), *SF3B1*^*mut*^ K700E mutated patients were more likely to have a lower CCSS scores (0/1) compared to non-K700E MDS patients [39 (80%) vs. 19 (53%); p=0·011].

### Therapy and outcomes

The majority of patients were treated with an HMA: 16/17 (94%) K700E mutated patients; 15/19 (79%) non-K700E mutated patients; and 217/277 (78%) *SF3B1*^*wt*^ patients. With a median follow-up of 15·6 months, *SF3B1*^*mut*^ MDS patients had a significantly better OS than *SF3B1*^*wt*^ patients. This was true in the entire cohort (not reached vs. 25·2 months, p=0·0003), low-grade MDS categories (not reached vs. 41·3 months, p=0·0015), and low-grade MDS-RS categories (not reached vs. 22·3 months, p=0·0004). When segregated based on *SF3B1* mutation types, 4 (7·3%) of the K700E mutated patients died, while 9 (23%; p=0·036) of non-K700E mutated patients died. The outcomes of K700E *SF3B1*^*mut*^ mutated MDS patients were significantly better compared to *SF3B1*^*wt*^ within the entire cohort (median OS, not reached vs. 25.2 months, p=0.0001), and within the low-grade MDS (median OS, not reached vs. 41.3 months, p=0·0015) and MDS-RS categories (median OS, not reached vs. 22.3 months, p=0.0001). In contrast, the outcomes of non-K700E *SF3B1*^*mut*^ MDS patients were similar to *SF3B1*^*wt*^ MDS patients, within the entire cohort (median OS, not reached vs. 25·2 months, p=0·2314), and within the low-grade MDS (median OS, not reached vs. 41·3 months, p=0·2598), and MDS-RS categories (median OS, not reached vs. 22·3 months, p=0·2327). Within the MDS-EB categories, compared to *SF3B1*^*wt*^, there were no significant differences in OS of K700E *SF3B1*^*mut*^ (median OS, not reached vs. 17·7 month, p=0·355) and non-K700E *SF3B1*^*mut*^ (median OS, 20·5 vs. 17·7 months; p=0·5477) patients (**Figure 3A-D**).

**Figure 3.**
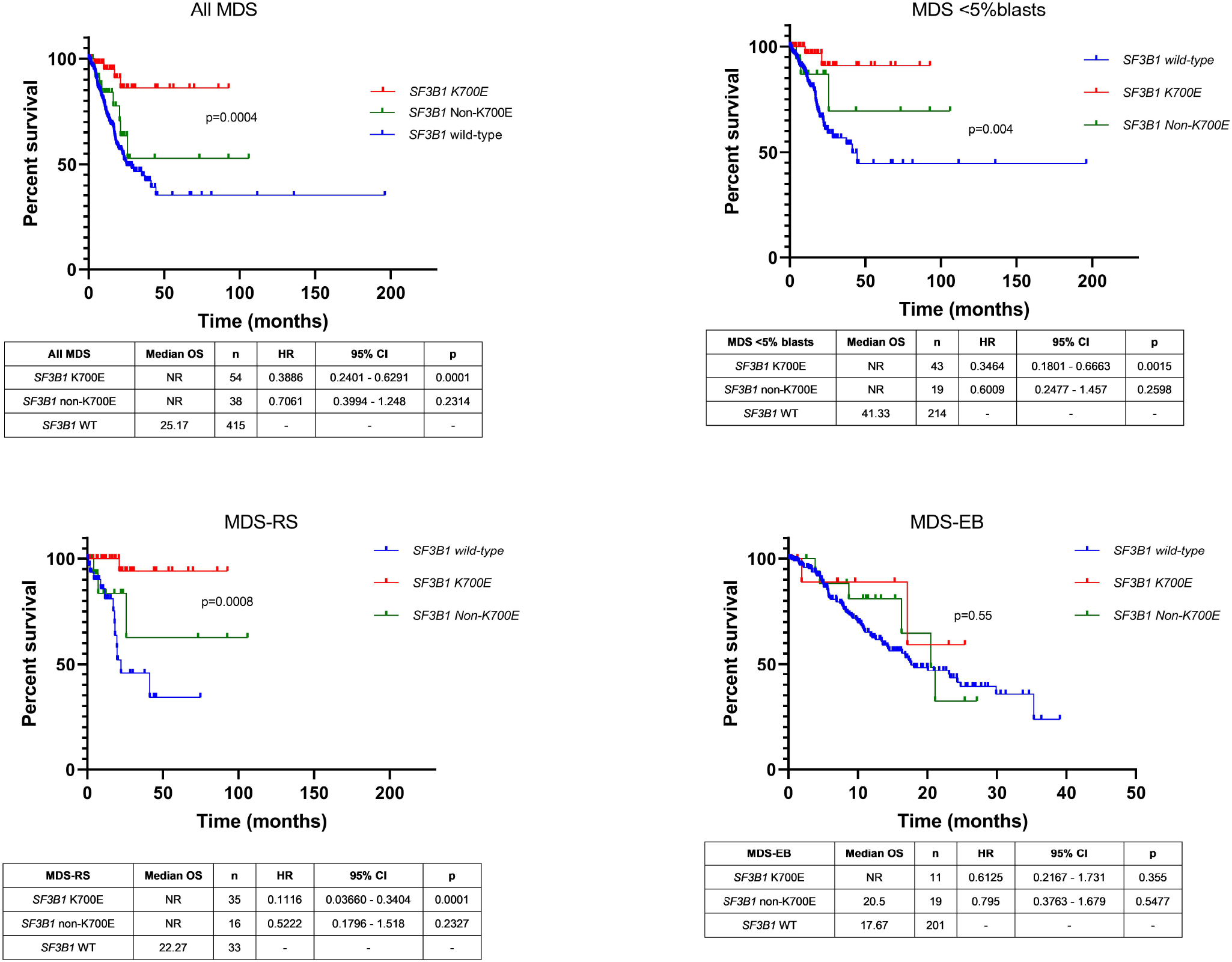
Overall survival (OS) of *SF3B1* mutated MDS subtypes compared to *SF3B1* wild-type MDS patients. OS of *SF3B1* mutated MDS patients was significantly better than wild-type patients. When segregated based on *SF3B1* mutation types, the outcome of *SF3B1* non-K700E patients was similar to wild-type MDS, within the entire cohort (A, median OS, not reached for both; p=0·02), within low-grade MDS (B), and MDS-RS (C) categories, while there were no significant differences in OS within MDS-EB categories (D). The outcome of *SF3B1* K700E patients was superior in all the categories. Kaplan-Meier curves show patients with *SF3B1* K700E MDS mutations (red), *SF3B1* non-K700E mutation MDS patients (green), and *SF3B1* wild-type MDS patients (blue).

Within the entire MDS cohort, by univariate analysis (**Table 3**), the following parameters were associated with worse outcomes: higher BM blasts percentage; lower hemoglobin, platelet and MCV; prior history of chemo-radiation (t-MDS); presence of CK; higher IPSS-R risk category; WHO MDS-EB category; absence of mutations in *SF3B1* K700E, *TET2*, and *U2AF1*; and presence of a *TP53* mutation. The outcome was superior when the *SF3B1* mutation VAF was higher [p=0·026; OR, 0·96 (0·924-0·995)]. Non-K700E *SF3B1* mutations did not associate with OS. By multivariable analysis (using a p-value cut-off of 0·200), lower hemoglobin, higher IPSS-R category, absence of *SF3B1* K700E, and presence of *TP53* mutation were independent predictors of worse OS. Within the MDS-RS categories (MDS-RS-SLD and MDS-RS-MLD), independent prognostic factors of worse OS included lower platelet count and presence of mutations in *SF3B1* (non-K700E), *ASXL1, SRSF2*, and *TP53* (**Table 4**).

**Table 3.** Univariate analysis for overall survival in all newly diagnosed untreated MDS patients.

|  | **p-value** | **OR** | **95% CI** |
| --- | --- | --- | --- |
| Age | 0.412 | 1.006 | 0.992-1.020 |
| Gender | 0.052 | 0.720 | 0.518-1.002 |
| **BM blasts** | <0.001 | 1.081 | 1.047-1.117 |
| ANC | 0.330 | 1.013 | 0.987-1.041 |
| **Hemoglobin** | <0.001 | 0.693 | 0.625-0.769 |
| **Platelet count** | <0.001 | 0.995 | 0.992-0.997 |
| **Mean Corpuscular** | <0.001 | 0.958 | 0.943-0.974 |
| **Therapy-related** | <0.001 | 2.134 | 1.539-2.959 |
| **R-IPSS** |  |  |  |
| Very Low | 0.199 | 0.266 | 0.035-2.011 |
| Low | - | - | - |
| Intermediate | 0.002 | 2.896 | 1.491-5.627 |
| High | <0.001 | 3.911 | 2.075-7.372 |
| Very High | <0.001 | 10.682 | 6.030- |
| **Complex karyotype** | <0.001 | 4.910 | 3.483-6.920 |
| MDS-EB | <0.001 | 2.167 | 1.494-3.144 |
| **SF3B1** |  |  |  |
| Wild type | - | - | - |
| **K700E** | 0.001 | 0.175 | 0.064-0.473 |
| Non-K700E | 0.224 | 0.657 | 0.334-1.293 |
| **SF3B1 VAF** | 0.026 | 0.959 | 0.924-0.995 |
| **NGS mutations (seen in >5% of cases)** |  |  |  |
| *ASXL1* | 0.817 | 0.950 | 0.612-1.473 |
| *BCOR* | 0.114 | 0.324 | 0.080-1.309 |
| *DNMT3A* | 0.650 | 0.896 | 0.558-1.438 |
| *IDH1/IDH2* | 0.127 | 0.498 | 0.204-1.218 |
| *RAS* | 0.434 | 0.700 | 0.287-1.709 |
| *RUNX1* | 0.760 | 0.920 | 0.539-1.570 |
| *SRSF2* | 0.499 | 0.832 | 0.487-1.420 |
| **TET2** | 0.002 | 0.469 | 0.289-0.760 |
| **TP53** | <0.001 | 5.158 | 3.699-7.193 |
| **U2AF1** | 0.023 | 0.437 | 0.214-0.891 |

**Table 4.** Multivariate analysis for overall survival in all newly diagnosed untreated MDS patients (p-value cutoff of 0.200 from univariate analysis) *SF3B1* mutations were segregated by subtype.

|  | **p-value** | **OR** | **95% CI** |
| --- | --- | --- | --- |
| **Hemoglobin** | <0.001 | 0.805 | 0.713-0.909 |
| **R-IPSS** |  |  |  |
| Very Low | 0.282 | 0.326 | 0.042-2.512 |
| Low | - | - | - |
|  | 0.016 | 2.288 | 1.166-4.487 |
| **High** | 0.010 | 2.436 | 1.237-4.798 |
| **Very High** | <0.001 | 4.143 | 2.076-8.467 |
| ***SF3B1*** |  |  |  |
| Wild type | - | - | - |
| **K700E** | 0.041 | 0.288 | 0.087-0.951 |
| Non- | 0.804 | 0.913 | 0.445-1.873 |
| *BCOR* | 0.154 | 0.234 | 0.032-1.724 |
| ***TP53*** | <0.001 | 2.374 | 1.539-3.662 |
| *U2AF1* | 0.082 | 0.513 | 0.242-1.089 |

Given the known independent prognostic influence of both *SF3B1* and *TP53* mutations, we assessed the co-mutation pattern and outcomes of *SF3B1* mutations with *TP53* mutations and CK. A total of 14 *SF3B1*^*mut*^ patients had concurrent *TP53* mutations [(n=9; median VAF 25·8 (range, 2-68%)] or CK (n=5). We divided the patients into 4 categories: (1) *SF3B1*^*wt*^ MDS without *TP53*^*mut*^ or CK (n=219); (2) *SF3B1*^*mut*^ MDS without *TP53*^*mut*^ or CK (n=71); (3) *SF3B1*^*mut*^ MDS with *TP53*^*mut*^ or CK (n=14); and (4) *SF3B1*^*wt*^ MDS with *TP53*^*mut*^ or CK (n=153). No survival differences were noted between patients with *SF3B1*^*mut*^ MDS with or without mutated *TP53* and/or CK (median OS, not reached) and patients with *SF3B1*^*wt*^ MDS without *TP53* mutations/CK (44·3 months). But MDS patients with *TP53* mutations or CK had a significantly worse outcomes (median OS, 12·9 months, HR 1·46, p=0·001) in the absence of *SF3B1* mutations. Similar findings were noted within the low-grade MDS and MDS-RS categories, where worse OS was associated with *TP53* mutations or CK when *SF3B1* was wild-type. This suggests that *SF3B1* mutation may negate the poor prognostic effect of *TP53* mutation or CK (**Figure 4**).

**Figure 4.**
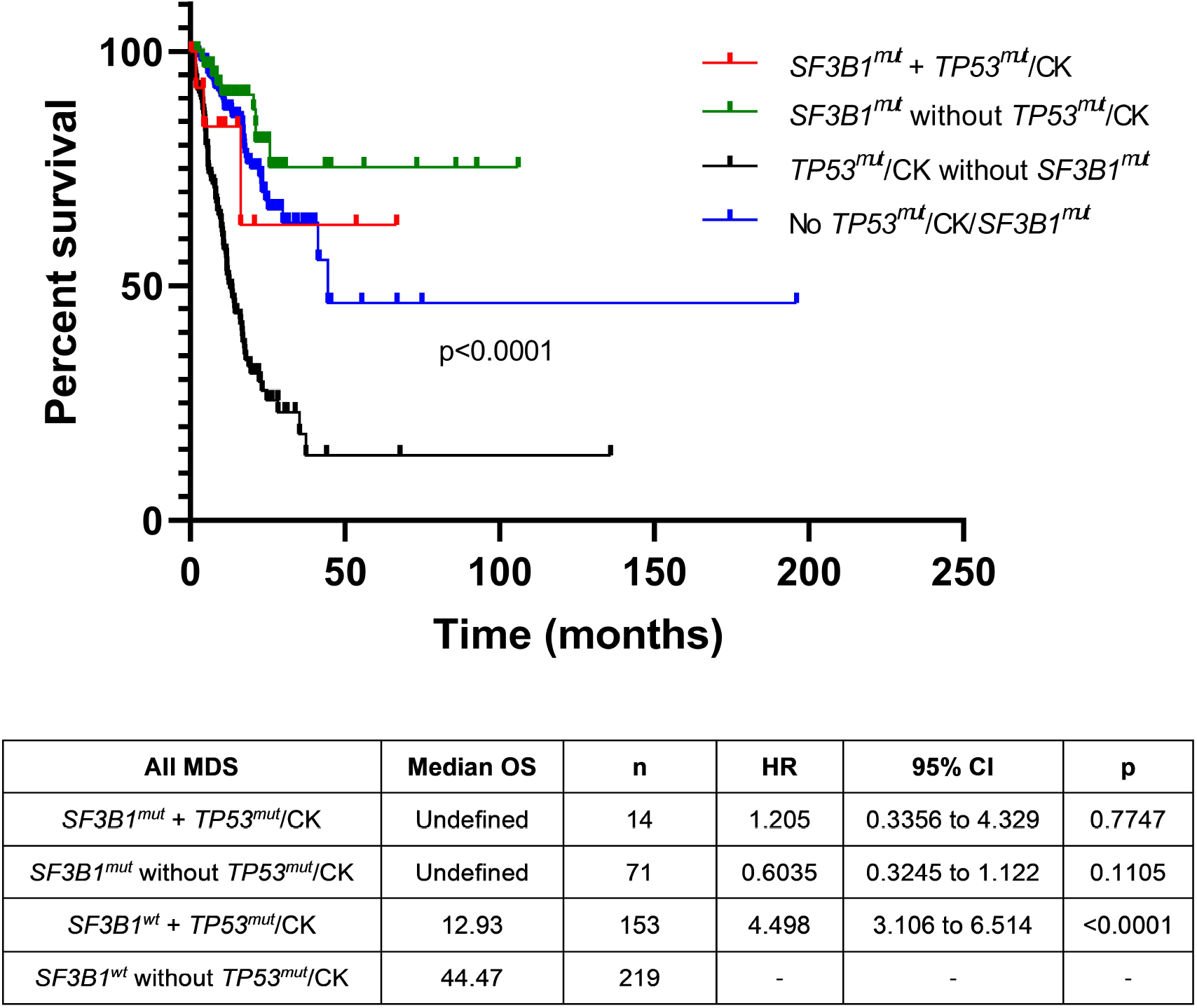
Survival differences in MDS based on concurrent *SF3B1* and *TP53* mutations and/or complex karyotype (CK). No overall survival differences were noted between *SF3B1* mutated MDS patients with or without mutated *TP53* and/or CK (median OS not reached) and *SF3B1* wild-type MDS patients without *TP53* mutations/CK (44·3 months). MDS patients with *TP53* mutations or CK had significantly worse outcomes (median OS 12·9 months, HR 1·46, p=0·001) in the absence of *SF3B1* mutations, suggesting that *SF3B1* mutation may negate the poor prognostic effects of *TP53* mutation or CK.

### *SF3B1* mutation acquired during disease course

*SF3B1* mutation is considered a founder clone, however we observed 2 patients in which the mutation arose during disease evolution. The first patient was a 74 year old man who was diagnosed with MDS-EB with trisomy 8 and mutations in *BCOR, DNMT3A* (x2), *EZH2, TET2*, and *U2AF1* (33% VAF). BM morphology did not show any ring sideroblasts. Flow cytometry analysis showed 5% B-cells with a CLL/SLL-like immunophenotype. Following HMA treatment, a 6-month follow-up BM aspirate analysis showed persistent MDS with 2% blasts and clonal evolution with additional sub-clonal mutations in *TET2* and *SF3B1* R625C (VAF, 29·6%), expansion in the *EZH2* clone, clearance of *BCOR*, and reduction in previously detected *TET2* and *U2AF1* mutations. As expected, given the acquisition of the *SF3B1* mutation, BM morphology showed 18% ring sideroblasts (**Figure 5A**). The second case was a 68 year old woman with pancytopenia. BM aspirate analysis showed MDS-EB with multi-lineage dysplasia and MF-3 fibrosis with an del(5q) abnormality in 16 of 20 metaphases. NGS analysis demonstrated a *JAK2* V617 (<5% VAF) mutation at baseline. The patient underwent treatment with an HMA and PD-1/CTLA-4 blockers, and transformed to AML over the next 4 months. Karyotype showed del(5q) abnormality. Additional mutations in *SF3B1* K700E (29% VAF) and *SETBP1* E862K (23·8% VAF) were acquired at transformation (**Figure 5B**). Both the patients underwent allogenic stem cell transplantation, and have remained in morphologic and molecular remission.

**Figure 5.**
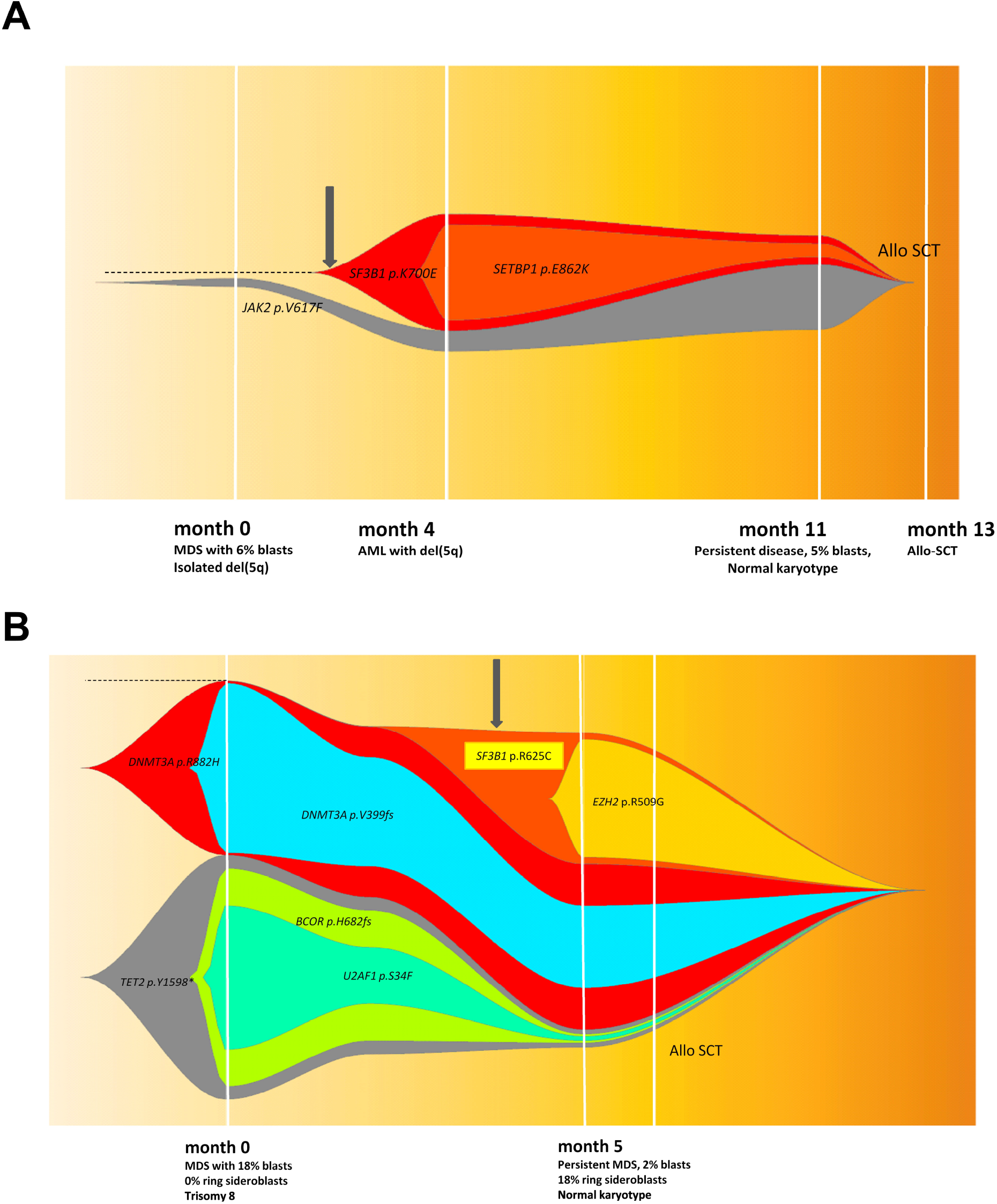
NGS analysis of serial of samples in two MDS patients who acquired *SF3B1* mutations over the course of the disease: in patient #1, at the time of AML progression (A), and in patient #2, post HMA therapy at the time of morphologically persistent disease with genotype change corresponding to phenotypic acquisition of ring sideroblasts (B). Representative FISH plots from both cases showing clonal architecture at different time points. In both cases, mutations cleared following allogeneic stem cell transplantation.

## DISCUSSION

The outcomes of *SF3B1*^*mut*^ MDS, although regarded as a favorable prognostic biomarker, are variable. In this study, to understand the heterogeneity in clinico-pathologic features and outcomes of different types of *SF3B1* mutations, we reviewed the entire spectrum of MDS, including low-grade (<5% blasts) through high-grade (≥5% blasts) MDS.

*SF3B1* mutations are observed in a third of MDS cases, are associated with ≥15% RS, and are overrepresented in the MDS-RS-SLD and MDS-RS-MLD WHO subtypes.^2,3,20^ Ring sideroblasts represent dysplastic erythroid precursors with abnormal iron-laden mitochondria encircling the nucleus.^21,22^ Based on the distinct gene expression profile of *SF3B1*^*mut*^ MDS and strong association with RS, the current WHO criteria classifies MDS with <5% BM blasts and 5-15% RS in the absence of isolated de(5q) with *SF3B1* mutation as equivalent to MDS-RS-SLD or MDS-RS-MLD characterized by ≥15% RS.^9^ *SF3B1* mutation is regarded as the only favorable prognostic biomarker in MDS.^2^ The most frequent hotspot *SF3B1* mutation is located on K700E, while the rest are located within exons 14-16.^3,4,14^

Consistent with the literature, our study found that *SF3B1*^*mut*^ MDS had favorable clinical characteristics, with enrichment in lower IPSS-R categories and ring sideroblastic WHO subtypes. However, when segregated by the type of mutation, we demonstrated that only a subset of *SF3B1*^*mut*^ MDS patients, those with K700E mutations, had favorable outcomes, while the outcomes of those with non-K700E mutations, noted in ∼40% of all *SF3B1* mutated patients, were similar to those with *SF3B1*^*wt*^ MDS. The only significant adverse clinical feature in non-K700E mutated *SF3B1* was lower ANC. Both K700E and non-K700E *SF3B1*^*mut*^ MDS patient groups showed frequent RS, with a median RS percentage exceeding 15% in both. Despite no significant difference in median BM blast percentage, these cases had a higher representation in the MDS-EB categories and a higher frequency of IPSS-R high cases. There were no karyotypic differences, but non-K700E *SF3B1*^*mut*^ MDS had a higher frequency of concomitant mutations in *RUNX1* (>25% cases) and *BCOR. RUNX1* mutation is an independent predictor of poor survival in MDS.^6,23^ Presence of K700E *SF3B1*^*mut*^ alone (not non-K700E mutations) was associated with favorable OS in both univariate and multivariate analysis, along with other prognostic factors including hemoglobin, IPSS-R category, and *TP53* mutations. The latter finding was true within low-grade MDS and MDS–RS categories. Thus, recognition of *SF3B1* mutation type by sequencing the entire coding region is important and has bearing on clinical management. Hence, incorporation of the type of *SF3B1* mutation rather than simply the presence of mutation and ring sideroblast percentage will be helpful for future WHO sub-classification.

*SF3B1* mutation is an MDS initiating clone arising in hematopoietic stem cells of lymphomyeloid origin, with the mutation often being the sole driver with a near-heterozygous VAF.^24,25^ Prior studies demonstrated that the presence of *SF3B1* mutation, in the context of clonal hematopoiesis or clonal cytopenia(s), was an independent predictor of progression to MDS, unlike isolated mutations in *DNMT3A* or *TET2*.^26^ Within our MDS cohort, >20% of *SF3B1*^*mut*^ MDS patients had common CHIP-associated gene mutations in *TET2* and *DNMT3A*. The presence of these additional gene mutations was not associated with multi-lineage dysplasia. *TP53* mutations were noted in >10% of *SF3B1*^*mut*^ MDS patients. Since *TP53* mutations associated with CK, as observed in ∼10% of MDS patients, we assessed the interplay of these 3 parameters on outcomes. Our findings showed that *SF3B1* mutations negate the poor prognostic effects of *TP53* mutations and/or CK. *SF3B1* mutation was less frequent within t-MDS, but was more frequent than *TP53* mutations within t-MDS with RS.

In addition to CK, other poor prognostic cytogenetic features in a subset of *SF3B1*^*mut*^ MDS included del(7q)/-7 and *EVI1/MECOM* rearrangement. Studies have shown enrichment of *SF3B1* mutations in AML with inv(3) or t(3;3) and overexpression of *EVI1* in *SF3B1*^*mut*^ MDS that transform to AML.^27,28^ In our cohort, the frequency of *EVI1* rearrangement was not significantly different from the wild-type cases. There were also 3 *SF3B1*^*mut*^ patients with t(1;3)(p36;q21), a rare but recurrent rearrangement in MDS/AML involving *PRDM16* (PR/SET domain containing 16) or *MEL1* (located on 1p36) and the enhancer element of *GATA2 (RPN1)*, located on 3q21. PRDM16 is zinc finger transcriptional regulator structurally similar to *MECOM* (*MDS1* and *EVI1* complex). Rearrangements of *PRDM16* in t(1;3)(p36;q21) and *MECOM* in inv(3)(q21q26.2) or t(3;3)(q21;q26.2) to the *GATA2/RPN1* locus lead to overexpression of the aberrant oncogenic short-forms of PRDM16s and EVI1 that lack the PR/SET domain. Aberrant Prdm16s has been implicated in leukemogenesis or progression by transformation of megakaryocyte-erythroid progenitors to myeloid leukemia stem cells.^29,30^

Within this single institution large dataset, we were able to evaluate *SF3B1* mutations in unique clinical scenarios, including isolated del(5q) (all of which were associated with K700E mutations), and t-MDS. A high proportion of cases with *TP53* mutation or CK (54%) were associated with t-MDS. Within *SF3B1*^*mut*^ MDS, a high proportion of t-MDS cases had concurrent *TP53* mutations and/or CK (57%), while fewer had *SF3B1* mutations without a *TP53* mutation or CK (19%). There were no differences in outcome based on therapy-related disease [HR 2·11, 95% CI 0·78-5·66, p=0·140].

Our study is a retrospective analysis of a large single-institutional series. Validation of these findings in an independent cohort, as well as prospective analysis of the outcome differences between K700E and non-K700E *SF3B1*^*mut*^ MDS patients, in the context of newer therapies such as luspatercept, is needed.

In summary, we show that ∼40% of *SF3B1*^*mut*^ MDS show non-K700E mutations. Hotspot K700E and non-K700E *SF3B1*^*mut*^ MDS show distinct clinical and mutational profiles, with K700E showing a significantly better OS compared to non-K700E and *SF3B1*^*wt*^. Only absence of *SF3B1*^*mut*^ K700E mutation independently predicted for worse OS in MDS. Hence, identification of the *SF3B1* mutation type is important for risk stratification.

## Author Contributions

**RK-S:** Concept and design; administrative support; provision of study materials and patients; data collection, analysis, interpretation; and manuscript writing and final approval. **GM-B, KS, EJ, CB-R, YW, ND, FD, MR, KP, LJM, HK, GG-M:** Collection and assembly of data; data analysis and interpretation; manuscript writing; and final approval of manuscript. **SP:** Provision of study materials; data collection, and final approval. **KAS:** Administrative support; data analysis and interpretation; and manuscript writing and final approval.

## Acknowledgements

This work was supported in part by grants from the Ladies Leukemia League (RK-S), the National Institutes of Health, National Cancer Institute University of Texas MD Anderson Cancer Center Support award CA016672 (all authors) and the University of Texas MD Anderson MDS/AML Moon Shot (YW, KAS, FD, HK, and GG-M).

## REFERENCES

1 Papaemmanuil, E. et al. Clinical and biological implications of driver mutations in myelodysplastic syndromes. Blood, The Journal of the American Society of Hematology 122, 3616–3627 (2013).

2 Malcovati, L. et al. SF3B1 mutation identifies a distinct subset of myelodysplastic syndrome with ring sideroblasts. Blood 126, 233–241, doi: 10.1182/blood-2015-03-633537 (2015).

3 Papaemmanuil, E. et al. Somatic SF3B1 mutation in myelodysplasia with ring sideroblasts. New England Journal of Medicine 365, 1384–1395 (2011).

4 Yoshida, K. et al. Frequent pathway mutations of splicing machinery in myelodysplasia. Nature 478, 64–69 (2011).

5 Chen, J. et al. Myelodysplastic syndrome progression to acute myeloid leukemia at the stem cell level. Nat Med 25, 103–110, doi: 10.1038/s41591-018-0267-4 (2019).

6 Malcovati, L. et al. SF3B1-mutant myelodysplastic syndrome as a distinct disease subtype-A Proposal of the International Working Group for the Prognosis of Myelodysplastic Syndromes (IWG-PM). Blood (2020).

7 Migdady, Y. et al. Clinical outcomes with ring sideroblasts and SF3B1 mutations in myelodysplastic syndromes: MDS clinical research consortium analysis. Clinical Lymphoma Myeloma and Leukemia 18, 528–532 (2018).

8 Montalban-Bravo, G. & Garcia-Manero, G. Myelodysplastic syndromes: 2018 update on diagnosis, risk-stratification and management. American journal of hematology 93, 129–147 (2018).

9 Arber, D. A. et al. The 2016 revision to the World Health Organization classification of myeloid neoplasms and acute leukemia. Blood 127, 2391–2405, doi: 10.1182/blood-2016-03-643544 (2016).

10 Kanagal-Shamanna, R. et al. Validation of the 2016 revisions to the WHO classification in lower-risk myelodysplastic syndrome. Am J Hematol 92, E168–E171, doi: 10.1002/ajh.24776 (2017).

11 Fenaux, P. et al. Luspatercept in Patients with Lower-Risk Myelodysplastic Syndromes. New England Journal of Medicine 382, 140–151 (2020).

12 Alsafadi, S. et al. Cancer-associated SF3B1 mutations affect alternative splicing by promoting alternative branchpoint usage. Nature communications 7, 1–12 (2016).

13 Darman, R. B. et al. Cancer-Associated SF3B1 Hotspot Mutations Induce Cryptic 3’ Splice Site Selection through Use of a Different Branch Point. Cell Rep 13, 1033–1045, doi: 10.1016/j.celrep.2015.09.053 (2015).

14 Obeng, E. A. et al. Physiologic expression of Sf3b1K700E causes impaired erythropoiesis, aberrant splicing, and sensitivity to therapeutic spliceosome modulation. Cancer cell 30, 404–417 (2016).

15 Dalton, W. B. et al. The K666N mutation in SF3B1 is associated with increased progression of MDS and distinct RNA splicing. Blood advances 4, 1192 (2020).

16 Arber, D. A. et al. The 2016 revision to the World Health Organization (WHO) classification of myeloid neoplasms and acute leukemia. Blood, doi: 10.1182/blood-2016-03-643544 (2016).

17 Greenberg, P. L. et al. Revised international prognostic scoring system for myelodysplastic syndromes. Blood 120, 2454–2465, doi: 10.1182/blood-2012-03-420489 (2012).

18 Kanagal-Shamanna, R. et al. Myeloid neoplasms with isolated isochromosome 17q demonstrate a high frequency of mutations in SETBP1, SRSF2, ASXL1 and NRAS. Oncotarget 7, 14251 (2016).

19 Cheson, B. D. et al. Clinical application and proposal for modification of the International Working Group (IWG) response criteria in myelodysplasia. Blood 108, 419–425, doi: 10.1182/blood-2005-10-4149 (2006).

20 Malcovati, L. & Cazzola, M. Recent advances in the understanding of myelodysplastic syndromes with ring sideroblasts. Br J Haematol 174, 847–858, doi: 10.1111/bjh.14215 (2016).

21 Della Porta, M. G. et al. Minimal morphological criteria for defining bone marrow dysplasia: a basis for clinical implementation of WHO classification of myelodysplastic syndromes. Leukemia 29, 66–75, doi: 10.1038/leu.2014.161 (2015).

22 Cazzola, M. & Invernizzi, R. (Haematologica, 2011).

23 Bejar, R. et al. Clinical effect of point mutations in myelodysplastic syndromes. New England Journal of Medicine 364, 2496–2506 (2011).

24 Mortera-Blanco, T. et al. SF3B1-initiating mutations in MDS-RSs target lymphomyeloid hematopoietic stem cells. Blood 130, 881–890 (2017).

25 Mian, S. A. et al. SF3B1 mutant MDS-initiating cells may arise from the haematopoietic stem cell compartment. Nature communications 6, 1–14 (2015).

26 Malcovati, L. et al. Clinical significance of somatic mutation in unexplained blood cytopenia. Blood 129, 3371–3378 (2017).

27 Shiozawa, Y. et al. Gene expression and risk of leukemic transformation in myelodysplasia. Blood, The Journal of the American Society of Hematology 130, 2642–2653 (2017).

28 Papaemmanuil, E. et al. Genomic classification and prognosis in acute myeloid leukemia. New England Journal of Medicine 374, 2209–2221 (2016).

29 Hu, T. et al. PRDM16s transforms megakaryocyte-erythroid progenitors into myeloid leukemia– initiating cells. blood 134, 614–625 (2019).

30 Mochizuki, N. et al. A novel gene, MEL1, mapped to 1p36. 3 is highly homologous to the MDS1/EVI1 gene and is transcriptionally activated in t (1; 3)(p36; q21)-positive leukemia cells. Blood, The Journal of the American Society of Hematology 96, 3209–3214 (2000).

